# Combinatorial genetic strategy accelerates the discovery of cancer genotype-phenotype associations

**DOI:** 10.1101/2023.04.12.536652

**Authors:** Shan Li, Alicia Wong, Huiyun Sun, Vipul Bhatia, Gerardo Javier, Sujata Jana, Robert B. Montgomery, Jonathan L. Wright, Hung-Ming Lam, Andrew C. Hsieh, Bishoy M. Faltas, Michael C. Haffner, John K. Lee

## Abstract

Available genetically-defined cancer models are limited in genotypic and phenotypic complexity and underrepresent the heterogeneity of human cancer. Herein, we describe a combinatorial genetic strategy applied to an organoid transformation assay to rapidly generate diverse, clinically relevant bladder and prostate cancer models. Importantly, the clonal architecture of the resultant tumors can be resolved using single-cell or spatially resolved next-generation sequencing to uncover polygenic drivers of cancer phenotypes.

## Main

Most cancers are not driven by a single oncogenic driver but are rather the sum of multiple genetic perturbations that occur during tumor evolution. However, the functional impact of most genomic abnormalities found in cancers remains largely unknown. Genetically-engineered mouse models are a standard approach to functionally define genetic drivers in cancer but they are costly, slow, and do not allow facile manipulation of more than a few genes. To address these limitations, we developed a novel methodology incorporating barcoded lentiviral (LV) libraries encoding cancer-associated genetic events into primary epithelial cells at a high multiplicity-of-infection (MOI) that are engrafted in mice for tumorigenic selection. This system enables LV barcode sequencing of tumors to identify cooperative oncogenic drivers of malignant transformation and specific cancer phenotypes.

Organotypic or organoid cultures permit the expansion of primary epithelial cells while maintaining their complex organization and tissue function. A major barrier to higher-order genetic studies in this context has been inefficient transgenesis using available LV transduction protocols. We hypothesized that enforced cell-virus contact in a constrained volume of gel matrix could increase LV transduction efficiency. Primary mouse bladder urothelial (mBU) and prostate epithelial (mPE) cells were isolated by fluorescence-activated cell sorting (FACS) based on a lineage-negative (Lin^−^: CD45^−^CD31^−^Ter119^−^), EpCAM^+^CD49f^high^ immunophenotype (Extended Data Fig. 1a), as these populations self-renew at high frequencies^1^ and readily establish organoids in culture (Extended Data Fig. 1b). Cells were mixed into cold Matrigel containing concentrated LV expressing green fluorescent protein (GFP) prior to seeding and polymerization of organoid droplets^2^. mBU and mPE cells could be completely transduced, delivering up to 10-20 copies per cell (Figs. 1a and 1b). We next developed a barcoding system to characterize the distribution of unique proviral copies per cell. LV constructs were barcoded with matching 10 nucleotide sequences distal to the 5’ long terminal repeat (LTR) and proximal to the 3’ LTR and produced as a pool. A custom single-cell amplicon panel was designed on the Mission Bio Tapestri platform to enable sensitive enumeration of multiple uniquely barcoded LVs per cell (Extended Data Fig. 1c). This approach was validated using a defined population of NIH 3T3 cells engineered with LVs to harbor up to 4 unique LV barcodes per cell (Extended Data Fig. 1d). mPE were transduced with a diverse barcoded LV pool at varying MOIs and single-cell amplicon sequencing showed relatively normal distributions of proviral copies per cell (Extended Data Fig. 1e and Fig. 1c).

**Fig. 1.**
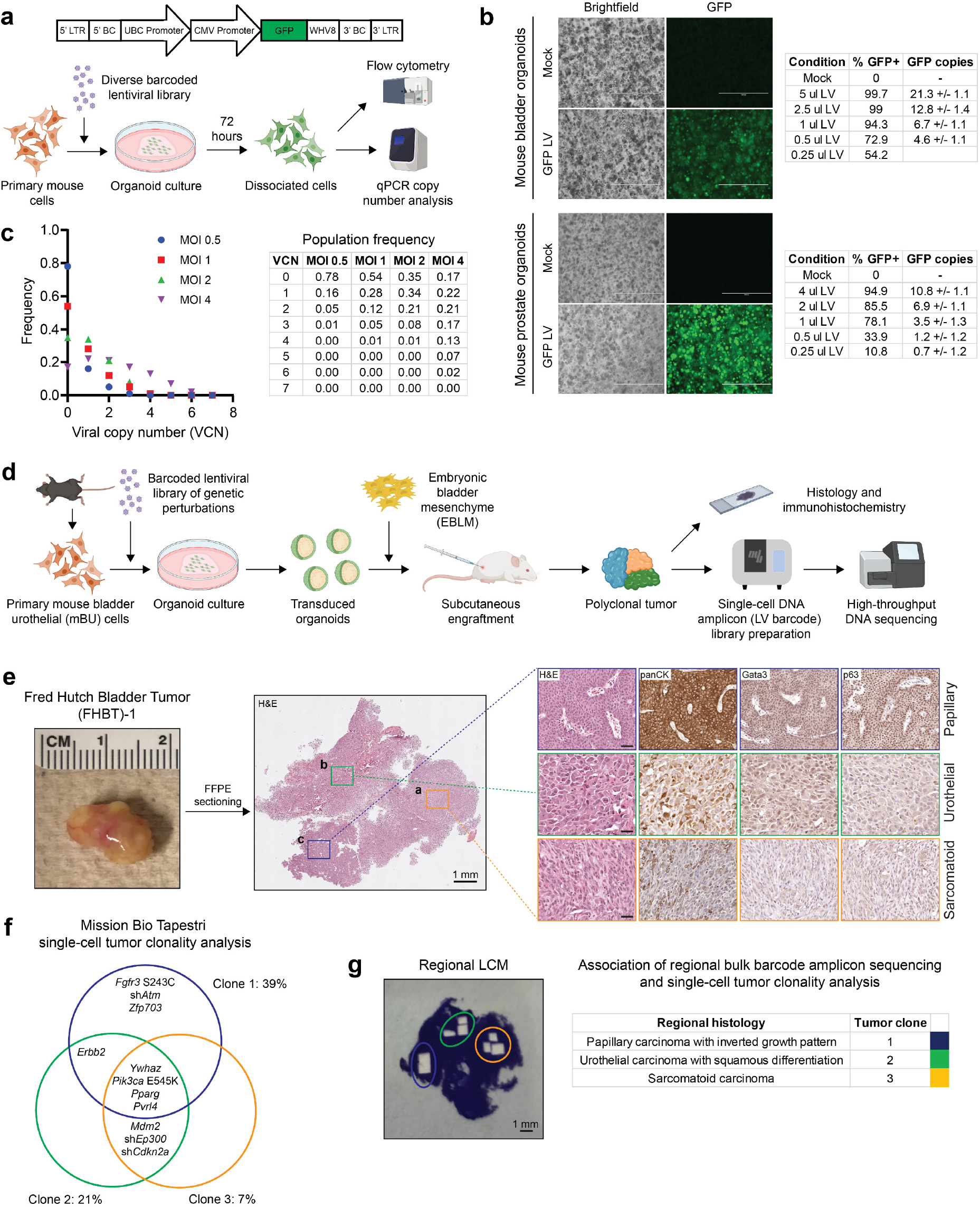
Efficient lentiviral transduction of primary epithelial cells at high multiplicity-of-infection and transformation of urothelial cells to tumors with mixed cancer histologies. (**a**) *Top*, Schematic of a LV construct with matching barcodes (BC) at the 5’ and 3’ ends. *Bottom*, Overview of experiments with LV infection of primary mouse cells in organoid culture and quantification of transduction. (**b**) *Left*, Brightfield and GFP images of mouse bladder or prostate organoids 72 hours after mock or GFP LV transduction. Scale bar=400 μm. *Right*, Tables summarizing quantification of LV transduction by flow cytometry and LV copies of GFP (+/− SD) by qPCR. (**c**) *Left*, Plot of the distribution of LV copies per mPE cell at different MOIs 72 hours after transduction. *Right*, Table summarizing VCN population frequencies at varying MOIs. (**d**) Scheme of the mBU organoid transformation assay to uncover functional genotype-phenotype associations in bladder cancer. (**e**) *Left*, Gross image of a tumor arising from mBU transformed with BU-LVp. *Middle*, low-magnification image of the H&E-stained tumor section. *Right*, high-magnification images of H&E- and IHC-stained sections of regions with distinct histologies. Scale bars=50 μm. (**f**) Clonal architecture of the three dominant clones in the tumor as determined by Mission Bio Tapestri single-cell analysis of LV barcodes. (**g**) *Left*, Tumor tissue section after LCM of the histologically distinct regions. *Right*, Table showing the associations between regional tumor histologies and clones in **f** based on LCM and bulk DNA amplicon sequencing of LV barcodes.

To determine the utility of this approach in understanding the initiation/progression of bladder and prostate cancer, we selected commonly mutated genes from cancer genome sequencing studies^3–5^ (Extended Data Fig. 2a) and cloned these as open reading frames (ORF) or short hairpin RNAs (shRNA) into barcoded LV constructs to mimic gain- or loss-of-function events (Extended Data Fig. 2b, Extended Data Table 1). At least three shRNA from The RNAi Consortium (TRC) targeting each gene were tested for knockdown in 3T3 cells by quantitative polymerase chain reaction. shRNA demonstrating the most potent knockdown of target gene expression was incorporated into the LV libraries (Extended Data Fig. 2c). A bladder urothelial LV pool (BU-LVp) of 33 genes and a prostate epithelial LV pool (PE-LVp) of 24 genes were produced in arrayed format to avoid LV barcode recombination and concentrated by ultracentrifugation (Extended Data Fig. 3a). Infectivity (representation) of each LV was evaluated by transducing either mBU or mPE cells with the respective LV pool and performing bulk amplicon sequencing of LV barcodes (Extended Data Fig. 3b). Initial LV pools demonstrated over 10-fold overrepresentation of shRNA vectors relative to ORF vectors (Extended Data Fig. 3c), presumably due to more efficient viral packaging because of the reduced length between LTRs of the transfer plasmid^6^. This data was applied to adjust the cell surface area of producer cells for subsequent arrayed LV library production, leading to near normalization of the representation of shRNA and ORF vectors (Extended Data Fig. 3d).

We adopted an approach in which primary mBU and mPE cells infected with BU-LVp or PE-LVp at high MOI in organoids were recombined with inductive mouse E16 embryonic bladder mesenchyme (EBLM)^7^ or urogenital sinus mesenchyme (UGSM)^8^ and subsequently grafted subcutaneously in NOD scid gamma (NSG) mice to enable biological selection for tumorigenic clones (Fig. 1d). A representative tumor derived from primary mBU cells transduced with BU-LVp exhibited three morphologically distinct regions consistent with papillary urothelial carcinoma with inverted growth pattern, urothelial carcinoma with squamous differentiation, and sarcomatoid urothelial carcinoma which were also supported by Gata3, p63, and pan-cytokeratin immunostaining (Fig. 1e). Single-cell DNA amplicon sequencing was performed to enumerate the LV barcodes for the determination of clonal architecture and deconvolution of LV-delivered genetic events putatively involved in tumorigenesis. Three major clones harboring distinguishable sets of LV barcodes were identified (Fig. 1f) but spatial resolution was lost due to single-cell dissociation. To associate histology with clonality, we performed laser capture microdissection (LCM) of the three regions on stained tissue sections and performed bulk DNA amplicon sequencing (Fig. 1g). The papillary urothelial carcinoma was uniquely associated with *Fgfr3* S243C, sh*Atm*, and *Zfp703*, in addition to the common *Ywhaz, Pik3ca* E545K, *Pparg*, and *Pvrl4* observed in all three dominant clones.

Cancer genomics studies have shown that activating mutations in *FGFR3* are highly enriched in papillary urothelial carcinomas^3,9^. We further validated these findings in the mouse urothelial transformation assay in independent experiments using a defined LV pool of *Fgfr3* S243C, *Ywhaz, Pik3ca* E545K, *Pparg*, and *Pvrl4* (Extended Data Fig. 4a) which produced tumors with papillary urothelial carcinoma with inverted growth pattern (Extended Data Fig. 4b).

Several tumors in the Fred Hutch Bladder Tumor (FHBT) series have been generated using this methodology including those with pure urothelial carcinoma and others with mixtures of histologic subtypes (Fig. 2a and Extended Data Fig. 5). The urothelial origin of these tumors was supported by GFP staining (Fig. 2b-d), which was positive even in regions of sarcomatoid carcinoma with low/absent pan-cytokeratin staining (Fig. 2b). We conducted molecular profiling of these tumors and their regional tumor histologies by LCM and RNA-seq analysis. Principal component analysis (PCA) of the gene expression data showed that squamous and sarcomatoid subtypes clustered together and were separate from urothelial and papillary urothelial carcinomas (Fig. 2e). The BASE47 subtype predictor^10^ generally called the tumors with papillary and papillary squamous subtypes as luminal and the squamous and sarcomatoid histologies as basal, consistent with an established relationship between sarcomatoid differentiation and the basal subtype^11^ (Fig. 2f). The Consensus Molecular Classifier^12^ revealed that the non-papillary urothelial histologies showed neuroendocrine (NE)-like gene expression with low/absent luminal and basal gene signatures (Fig. 2g and Extended Data Fig. 6a). Gene set enrichment analysis (GSEA) from comparing these tumor histologies in a pairwise manner revealed enrichment of genes associated with epithelial-to-mesenchymal transition in sarcomatoid carcinoma, as expected from prior molecular analyses of human tumors^11^ (Fig. 2h). We further confirmed the relevance of our FHBT models by projecting their RNA expression patterns onto PCA plots of human muscle-invasive bladder cancers from the TCGA-BLCA cohort^3^ (Fig. 2i) and established N-butyl-N-(4-hydroxybutyl)-nitrosamine (BBN)-induced mouse bladder cancer models^13^ (Extended Fig. 6b) to show that they occupy overlapping space.

**Fig. 2.**
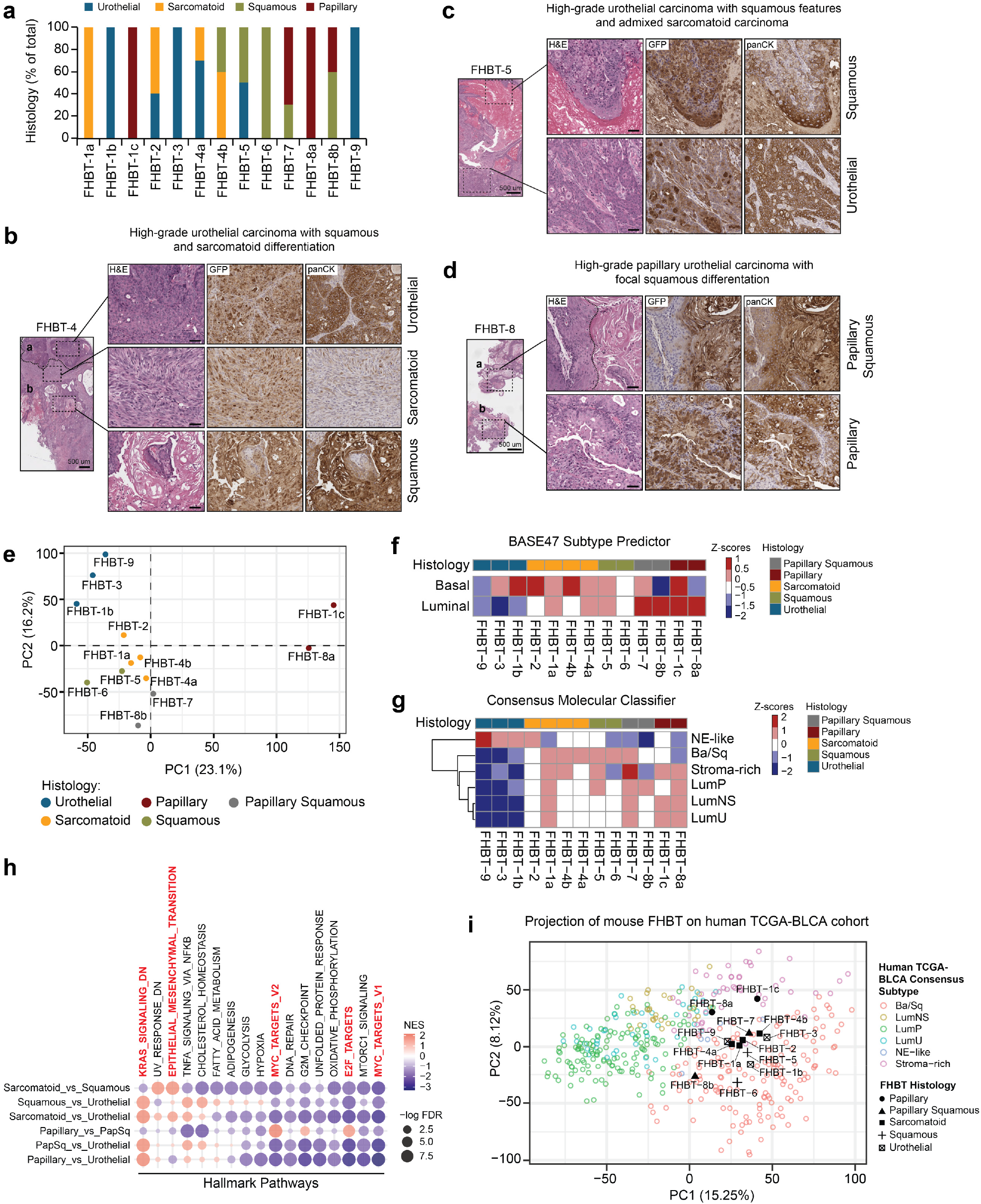
Rapid generation of a series of clinically relevant and phenotypically diverse bladder cancer models. (**a**) Bar graph showing the representation of cancer histologies present across a series of tumors (FHBT) generated using mBU transformed with BU-LVp. (**b**-**d**) Low- and high-magnification images of H&E-stained sections and high-magnification images of IHC-stained sections for GFP and pan-cytokeratin (panCK) expression depicting mixed histologies present within the same tumor. Scale bars=50 μm. (**e**) PCA plot showing FHBT series color-coded based on histology. Heatmaps showing the histologies of the FHBT series relative to (**f**) basal and luminal signature scores for the BASE47 subtype predictor and (**g**) signature scores for the Consensus Molecular Classifier. (**h**) Pre-ranked GSEA dotplot of Hallmark Pathways based on differentially expressed genes (FDR <0.001) in pairwise histology comparisons. (**i**) PCA projection plot of FHBT samples over the TCGA-BLCA samples color-coded by Consensus Molecular Classification.

mPE cells transduced with PE-LVp and engrafted in mice (Extended Data Fig. 7a) also gave rise to mixed cancer morphologies. One tumor showed high-grade prostate adenocarcinoma with focal pleomorphic giant cells (Extended Data Fig. 7b), a rare histologic subtype associated with poor prognosis^14^ that may contribute to therapeutic resistance and lethality^15^. Immunostaining revealed Hoxb13 and Ar expression in both histologies with pronounced nuclear p53 expression in the pleomorphic giant cells (Extended Data Fig. 7b). We isolated large (pleomorphic giant cell carcinoma) and small (adenocarcinoma) cells from dissociated tumors using a flow cytometry-based strategy, propagated these cells briefly in organoid cultures, then dissociated the cells and stained with the nuclear dye Hoechst 33342 to further isolate cells based on DNA content for downstream single-cell LV barcode enumeration (Extended Data Fig 7c). This single-cell clonality analysis revealed striking enrichment of sh*Kmt2c* in the putative pleomorphic giant cell clones (Extended Data Fig. 7d). Recent studies have established that KMT2C mediates DNA damage response in cancer^16,17^ and DNA damage repair alterations are common in human prostate adenocarcinoma with pleomorphic giant cell features^18^.

In sum, we describe a novel set of technologies that form a functional *in vivo* cancer genomics assay with efficient delivery of compound genetic perturbations from barcoded LV libraries and single-cell sequencing to rapidly investigate genotype-phenotype relationships in cancer initiation/progression from primary epithelial cells. These initial studies provide proof-of-principle of this approach but additional optimization is ongoing to enable the high-throughput study of tumorigenesis in the native tissue microenvironment in immune-competent hosts and more facile spatial association of genotype-phenotype.

## Methods

### Lentiviral constructs and lentiviral library production

Double-barcoded lentiviral vectors were generated from FU-CGW by sequentially inserting matched 10-nucleotide barcodes into the PacI site distal to the HIV FLAP using the Quick Ligation Kit (New England Biolabs) and PCR amplification of the WPRE sequence and barcode with insertion into the ClaI sites proximal to the 3’ LTR by HiFi DNA Assembly (New England Biolabs). ORFs were cloned into the EcoRI site of the double-barcoded lentiviral vectors by HiFi DNA Assembly. To generate shRNA lentiviral vectors, the Ubiquitin C promoter sequence was excised from the double-barcoded plasmid by digesting with PspXI and EcoRI. U6 promoter and shRNA cassettes were isolated by digesting pLKO.1 TRC shRNA clones with PspXI and EcoRI and were inserted into the digested double-barcoded plasmid using the Quick Ligation Kit. Individual lentiviruses were generated in arrayed format in 293T cells by co-transfection of each double-barcoded lentiviral ORF or shRNA plasmid with the helper plasmids pVSV-G, pMDL, and pRev using FuGENE HD Transfection Reagent (Promega). Lentiviral supernatants were collected 36 hours after transfection, pooled, and concentrated by ultracentrifugation in V-bottom polypropylene centrifuge tubes on a SW 32 Ti in an Optima XE 90 (Beckman Coulter) at a speed of 22,000 rpm at 4°C for 2 hours. Supernatants were aspirated and lentiviral pellets were resuspended in residual media and cryopreserved.

### shRNA screening

The top three to five shRNA sequences identified from The RNAi Consortium for each target gene were identified from the Genetic Perturbation Platform Web Portal at the Broad Institute. shRNA sequences were cloned into pLKO.1. pLKO.1-TRC control and pLKO.1-shRNA lentiviruses were generated and used to transduce 3T3 cells. 72 hours after lentiviral transduction, 3T3 cells were collected and RNA harvested using a RNeasy Mini Kit (Qiagen). Reverse transcription of RNA was performed using SuperScript IV Reverse Transcriptase (Invitrogen) per manufacturer’s instructions. qPCR was performed on a QuantStudio 6 using SYBR Green qPCR Master Mix (ThermoFisher Scientific) and primers specific to each target gene and *Ubc* as a control. All primers used for these studies are listed in Extended Data Table 2. Relative expression was calculated using ddCT analysis.

### Embryonic bladder mesenchyme and urogenital sinus mesenchyme preparation

Urogenital sinus mesenchyme was isolated and propagated as previously described^8^. E16 fetal bladders were also collected at the same time as the urogenital sinus and subjected to similar steps for preparation of embryonic bladder mesenchyme. Urogenital sinus mesenchyme and urogenital sinus mesenchyme were passaged less than five times prior to use in engraftment studies.

### Mouse bladder and prostate dissociation and organoid culture

Bladder and prostates from eight-to twelve-week-old male C57BL/6 mice (The Jackson Laboratory) were dissected and enzymatically dissociated in Collagenase/Dispase 1 mg/ml solution in DMEM media for 1 hour at 37ºC with continuous rocking. Cells were pelleted and supernatant aspirated. Trypsin-EDTA 0.25% solution was added to the cell pellet and incubated for 5 minutes at 37ºC. Trypsin was neutralized and cells were washed in DMEM + 10% FBS before filtering through a 40 μm cell strainer. Cells were stained with antibodies for fluorescence-activated cell sorting on a Sony SH800 Cell Sorter with collection of Lin^−^CD49f^high^EpCAM^high^ cells. 1-2 × 10^4^ bladder urothelial and prostate epithelial cells were resuspended in a total of 15 μl of growth factor-reduced Matrigel (Corning) +/− concentrated lentivirus and seeded as droplets in each 48-well tissue culture plate well. Cells were cultured as previously described^2^. Mouse bladder organoid culture media consisted of Advanced DMEM-F12, 10 mM HEPES, 2 mM GlutaMAX, B27 supplement, 1.25 mM N-acetylcysteine, 50 ng/ml hEGF, 100 ng/ml hNoggin and 500 ng/ml hR-spondin, 200 nM A83-01, and 10 μM Y-27632. Mouse prostate organoid culture media consisted of mouse bladder organoid culture media with the addition of 1 nM dihydrotestosterone.

### Organoid transformation assay

After 5-7 days of culture, transduced mouse bladder urothelial or prostate epithelial organoids were liberated by dissociating Matrigel matrix with Dispase 5 U/ml (STEMCELL Technologies). Organoids were washed with PBS and resuspended in ice cold Matrigel with either 10^5^ EBLM or UGSM and subcutaneously injected into the flanks of six-to eight-week-old male NSG (NOD-SCID-IL2Ry-null) mice (The Jackson Laboratory). For prostate epithelial transformation studies, mice were supplemented with testosterone through the subcutaneous implantation of 90-day release testosterone pellets (Innovative Research of America). Tumors were harvested when they reached 1 cm in maximal diameter. All animal care and studies were performed in accordance with an approved Fred Hutchinson Cancer Center Institutional Animal Care and Use Committee protocol and Comparative Medicine regulations.

### Copy number assay

DNA was extracted from organoids using a GeneJET Genomic DNA Purification Kit (ThermoFisher Scientific). Copy number analysis was performed by TaqMan Real-Time PCR Assay (ThermoFisher Scientific) using the TaqMan Copy Number Reference Assay, mouse, Tfrc (4458366) and EGFP TaqMan Copy Number Assay (Mr00660654_cn) on a QuantStudio 6. Genomic DNA extracted from the tails of transgenic C57BL/6 mice with one or two copies of GFP was used as a calibrator sample. GFP copy number was determined using ddCT analysis where sample copy number = calibrator copy number x 2^-ddCT^.

### Single-cell DNA amplicon sequencing library preparation and sequencing

A custom panel was designed for the Mission Bio Tapestri to amplify segments of ten mouse genes at two exons each, the 5’ and 3’ lentiviral barcodes, and lentiviral GFP. Libraries were generated either from cryopreserved or freshly dissociated tumor cells using the Mission Bio Tapestri Single-cell DNA Custom Kit according to the manufacturer’s recommendations. Single cells (3,000 to 4,000 cells per μl) were resuspended in Tapestri cell buffer and encapsulated using a Tapestri microfluidics cartridge, lysed, and barcoded. Barcoded samples were subjected to targted PCR amplification and PCR products were removed from individual droplets, purified with KAPA Pure Beads (Roche Molecular Systems), and used as a template for PCR to incorporate Illumina P7 indices. PCR products were purified by KAPA Pure Beads, and quantified by Qubit dsDNA High Sensitivity Assay (ThermoFisher Scientific). Sample quality was assessed by Agilent TapeStation analysis. Libraries were pooled and sequenced on an Illumina MiSeq or HiSeq 2500 with 150 bp paired-end reads in the Fred Hutchinson Cancer Center Genomics Shared Resource.

### Laser capture microdissection and DNA/RNA isolation for high-throughput sequencing

10 μm thick sections were cut from FFPE tumor tissues blocks and mounted onto PEN membrane frame slides (ThermoFisher Scientific). Sections were fixed with 95% ethanol for 1 minute, stained with 3% cresyl violet, and dehydrated through graded alcohols and xylene. Histology review and annotation was performed by a pathologist. Laser capture microdissection was performed on an Arcturus XT Laser Capture Microdissection System (ThermoFisher Scientific). Microdissected specimens were collected for DNA and RNA extraction. DNA was extracted using a GeneRead DNA FFPE Kit (Qiagen) and RNA was extracted using a RNeasy FFPE Kit (Qiagen) according to manufacturer’s protocols. Two-step PCR for amplification and sequencing library adaptor ligation was performed. The first PCR reaction consisted of 2x KAPA HiFi HotStart ReadyMix, 100 nM of 1º FWD primer (5’-TCGTCGGCAGCGTCAGATGTGTATAAGAGACAGCAAAATTTTCGGGTTTATTACAGG-3’), 100 nM of 1º REV primer (5’-GTCTCGTGGGCTCGGAGATGTGTATAAGAGACAGGCCGCTCGAGGACTATTAAG-3’), and 80 ng of genomic DNA. Thermal cycling conditions were 95ºC for 3 minutes; (95ºC for 30 seconds, 64ºC x 30 seconds, 72ºC x 30 seconds) x 25 cycles; 72ºC for 5 minutes, and hold at 4ºC. PCR cleanup was conducted using the Wizard SV Gel and PCR Clean-Up System (Promega) with elution in 30 μl of double distilled water. The second PCR reaction consisted of 2x KAPA HiFi HotStart ReadyMix, 140 nM of 2º i7 primer, 140 nM of 2º i5 primer, and 5 μl of elution from the PCR cleanup of the 1º PCR. Thermal cycling conditions were 95ºC for 3 minutes, (95ºC for 30 seconds, 61ºC x 30 seconds, 72ºC x 30 seconds) x 8 cycles; 72ºC for 5 minutes, and hold at 4ºC. The sequences of 2º primers used to incorporate dual-indexed Illumina sequencing adapters are displayed in Extended Data Table 3. PCR cleanup was conducted using the Wizard SV Gel and PCR Clean-Up System with elution in 30 μl of double distilled water. Sample quality was assessed by Agilent TapeStation analysis. Sequencing was performed on an Illumina MiSeq or HiSeq 2500 instrument using 150 bp single-end reads. PhiX sequences were excluded from the sequencing reads by Bowtie 2 v2.4.4^19^. Cutadapt v4.1^20^ was used to trim the reads to the barcode region. Then the trimmed reads were aligned to custom DNA references containing all barcodes using Bowtie 2. Samtools v1.11^21^ was used to extract read counts for each barcode. The RNA-seq libraries were prepared using a SMARTer Stranded Total RNA-Seq Kit v3 - Pico Input Mammalian (Takara Bio) and sequenced on an Illumina NovaSeq 6000 using a NovaSeq S4 flow cell with 100 bp paired end reads by MedGenome, Inc. Sequencing reads were mapped to mouse genome reference GRCm39 and gene expression was quantified and normalized using the UC Santa Cruz Computational Genomics Lab Toil RNA-seq pipeline v4.1.2^22^.

### Transcriptional subtype analysis and PCA projections

All computational analyses were carried out in RStudio Version 4.1.0. Mouse Ensembl genes were converted to Mouse Genome Informatics (MGI) gene symbols using the biomaRt package (https://bioconductor.org/packages/release/bioc/html/biomaRt.html). MGI gene symbols were then converted to their human orthologs by referencing the mouse-human ortholog database available from The Jackson Laboratory (http://www.informatics.jax.org/downloads/reports/HOM_MouseHumanSequence.rpt). The human ortholog matrix was used for downstream analysis in transcriptional subtype analysis. FHBT samples were classified using the BASE47 subtype predictor gene list^10^ and the ConsensusMIBC package v1.1^12^. Z-score means of genes and signature scores were calculated for each sample. Heatmaps of both the BASE47 and ConsensusMIBC results were generated using the pheatmap package v1.0.12 (https://www.rdocu-mentation.org/packages/pheatmap/versions/1.0.12/topics/pheatmap). For PCA analysis, the FPKM human ortholog matrix was normalized by log_2_ +1 transformation before performing mean-centered PCA using the prcomp package v3.6.2 (https://www.rdocumentation.org/pack-ages/stats/versions/3.6.2/topics/prcomp). Visualization of the PCA plot was performed using the factoextra package v1.0.7 (https://cran.r-project.org/web/packages/factoextra/index.html) and ggpubr package v0.6.0 (https://www.rdocumentation.org/packages/ggpubr/versions/0.6.0).

For PCA projections, RNA-seq count data from the FHBT, GSE220999, and TCGA-BLCA datasets were transformed to CPM and normalized to compare across each dataset using the DGEobj.utils package v1.0.6 (https://rdrr.io/cran/DGEobj.utils/). PCA projection of the FHBT data onto the TCGA-BLCA space was done by first generating a PCA of the TCGA-BLCA samples from the common genes between the FHBT and GSE220999 data. A PCA for both the FHBT and GSE220999 samples was then scaled by the eigenvalues of the TCGA-BLCA using the base package v3.6.2 (https://www.rdocumentation.org/packages/base/versions/3.6.2/topics/scale). A plot was constructed overlaying the reference TCGA-BLCA samples with either FHBT or GSE220999 tumor projections using ggplot2 v3.4.1 (https://cran.r-project.org/web/pack-ages/ggplot2/index.html). TCGA-BLCA samples were colored by their Consensus Molecular Classifier subtype. Differential gene expression analysis was performed pairwise between FHBT histologies using the DESeq2 package v1.38.3^23^. P-values were generated by Wald test and P-adj by Benjimini-Hochberg correction. Pre-ranked Gene Set Enrichment Analysis (Broad Institute) was conducted by inputting a ranked list of differentially expressed genes based on log_10_ transformed p-values from the DESeq2 analysis for each pairwise comparison. Dot plots were generated by plotting the Normalized Enrichment score and log transformed FDR for each pre-ranked GSEA output using ggplot2.

### Single-cell lentiviral barcode enumeration and clonality analysis

Raw sequencing reads were trimmed to the amplicon regions using the awk command. Barcode sequences in the reads were filtered and extracted using UMI-tools v1.0.0^24^. Processed reads were aligned to custom references containing all amplicon sequences using bwa-mem v0.7.17-r1188^25^. Samtools was used to extract amplicon counts for each barcode. Mouse cells with no GFP amplicon counts were removed. Counts per cell were normalized to total counts for each barcode. A minimum threshold normalized count of 1% of total counts was used to define the presence of a barcode in a cell. The clonal architecture of cells was determined by enumerating all cells containing each distinct combination of barcodes.

### Immunohistochemistry

Tumor samples were formalin-fixed and paraffin-embedded, sectioned to 5 μm thickness, and placed on positively charged glass slides. For each tumor, slides were stained with a standard H&E protocol. For immunohistochemical studies, tissue sections were deparaffinized in xylene and rehydrated in graded alcohols. Antigen retrieval was performed in pH 6.0 citrate buffer boiled for 30 minutes in a pressure cooker heated by microwave. Slides were washed with PBS, blocked with 2.5% horse serum for 30 minutes at room temperature, and then incubated with primary antibodies overnight at 4°C. Vector Laboratories ImmPRESS-HRP anti-rabbit or anti-mouse IgG (peroxidase) polymer detection kits were used to detect proteins. Liquid DAB+ Kit (Agilent Technologies) was used for chromogenic visualization of immunostaining. All sections were counter-stained with hematoxylin, dehydrated, and mounted with a glass coverslip.

### Antibodies

Antibodies used for FACS: Human/mouse/bovine integrin alpha 6/CD49f PE-conjugated antibody (FAB13501P, R&D Systems); PE/Cyanine 7 anti-mouse CD325 (Ep-CAM) antibody (118216, BioLegend); CD31 (PECAM-1) monoclonal antibody (390), FITC (11-0311-82,eBioscience); CD45 monoclonal anitbody (30-F11), FITC (11-0451-85, eBioscience); TER-119 monoclonal antibody (TER-119), FITC (11-5921-82, eBioscience). Antibodies used for immunohistochemistry: Rabbit polyclonal panCK (ab9377, Abcam, 1:100); rabbit monoclonal GFP antibody (clone D5.1, Cell Signaling, 1:100); rabbit polyclonal p63 antibody (12143-1-AP, Proteintech, 1:200); mouse monoclonal p53 antibody (clone 1C12, Cell Signaling, 1:500); rabbit monoclonal HOXB13 antibody (clone D7N8O, Cell Signaling, 1:50); rabbit polyclonal AR antibody (06-680, Millipore, 1:2,000); rabbit monoclonal GATA3 antibody (clone D13C9, Cell Signaling, 1:200); rabbit monoclonal CD44 antibody (clone E7K2Y, Cell Signaling, 1:100).

### Statistical analyses

Data analysis was performed on GraphPad Prism 9 (GraphPad Software, Inc.). qPCR results were analyzed in Excel. Statistical significance was determined using the unpaired two-tailed Student t test. Results are depicted as mean + SD unless stated otherwise. For all statistical tests, p-values of <0.05 were considered significant.

## Acknowledgements

We thank Colm Morrissey, Li Xin, and Peter Nelson for critical discussion and review of this work. This work was supported by NIH DP2 CA271301 (J.K.L.), R01 CA276308 (A.C.H.), R37 CA230617 (A.C.H.), Bladder Cancer Advocacy Network Research Innovator Award (J.K.L.), Department of Defense Peer Reviewed Cancer Research Program Career Development Award W81XWH-19-1-0569 (J.K.L.), Department of Defense Prostate Cancer Research Program Early Investigator Research Award W81XWH-20-1-0083 (S.L.), and Department of Defense Peer Reviewed Cancer Research Program Horizon Award W81XWH-19-1-0658 (S.J.). We acknowledge support from the Seattle Translational Tumor Research Program in Bladder Cancer and the University of Washington Medicine Urethral Cancer Research Fund provided by donor Don L. Rich. This research was also supported by the Flow Cytometry, Experimental Histopathology, and Genomics Shared Resources of the Fred Hutch/University of Washington Cancer Consortium funded by NIH P30 CA015704.

