## Extended Data Figures and Tables for "Combinatorial genetic strategy accelerates the discovery of cancer genotype-phenotype associations"

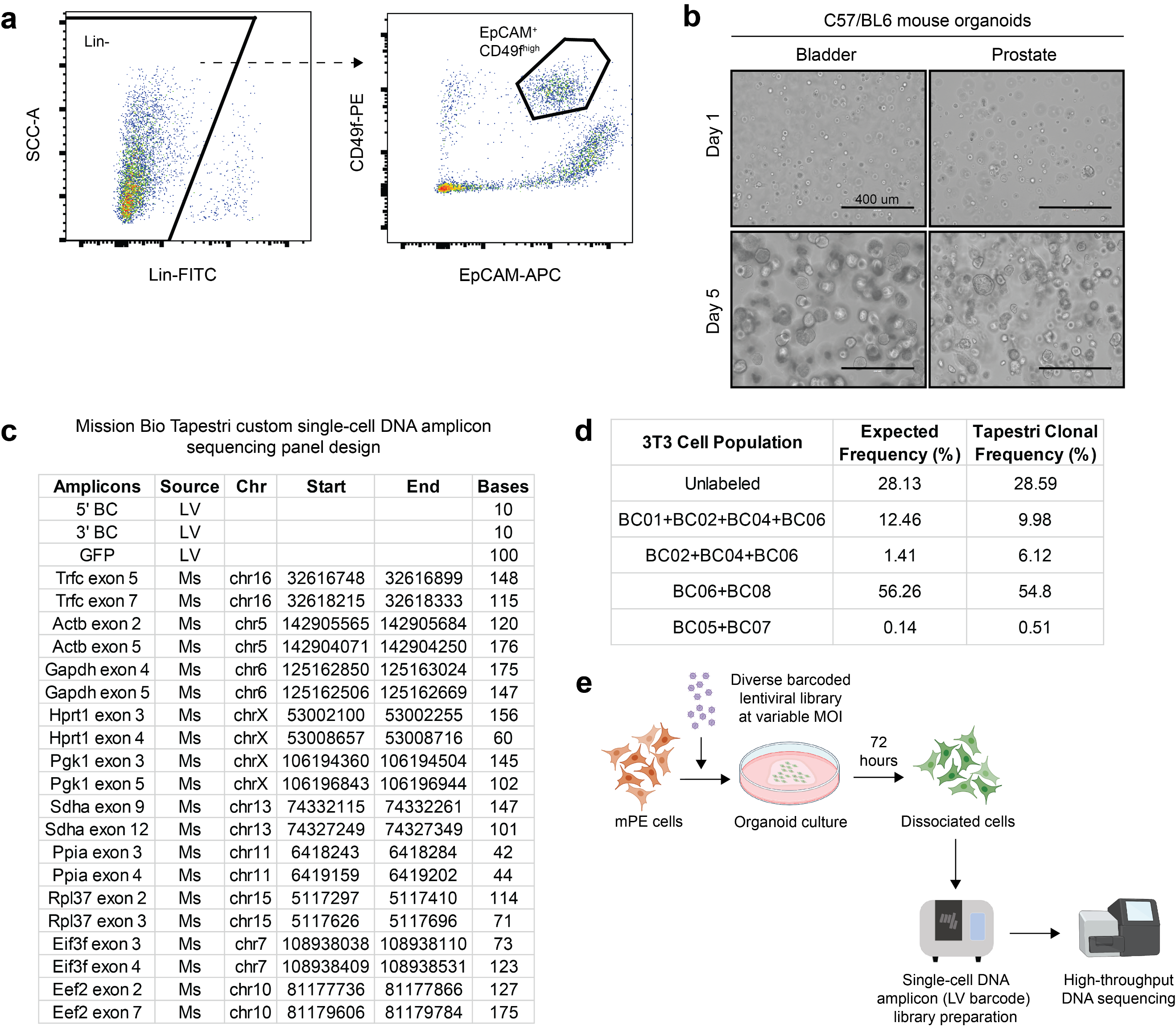

**Extended Data Fig. 1. Isolation of mouse bladder urothelial and prostate epithelial cells for organoid culture and design/validation of a custom Mission Bio Tapestri single-cell DNA amplicon sequencing panel.** (**a**) Representative flow cytometry plot for the isolation of mouse bladder urothelial and prostate epithelial from dissociated tissues based on a Lin^-^(CD45^-^CD31^-^Ter119^-^) EpCAM^+^CD49f^high^ immunophenotype. (**b**) Images of organoid cultures of mouse bladder urothelial and prostate epithelial cells on day 1 and day 5 after seeding. (**c**) Table showing the amplicons represented in a custom Mission Bio Tapestri single-cell DNA amplicon sequencing panel. (**d**) Table showing results of a validation study where a defined mixture of 3T3 cells with an unlabeled population and others labeled with combinations of lentiviruses encoding distinct barcodes were analyzed using the Mission Bio Tapestri single-cell DNA amplicon sequencing panel to determine clonality. ~2,000 cells were analyzed. (**e**) Overview of experiments with infection of mouse prostate epithelial (mPE) cells with a diverse barcoded lentiviral library in organoid culture across a range of multiplicity-of-infection (MOI) and quantification of viral copy number per cell across the population by single-cell amplicon sequencing.

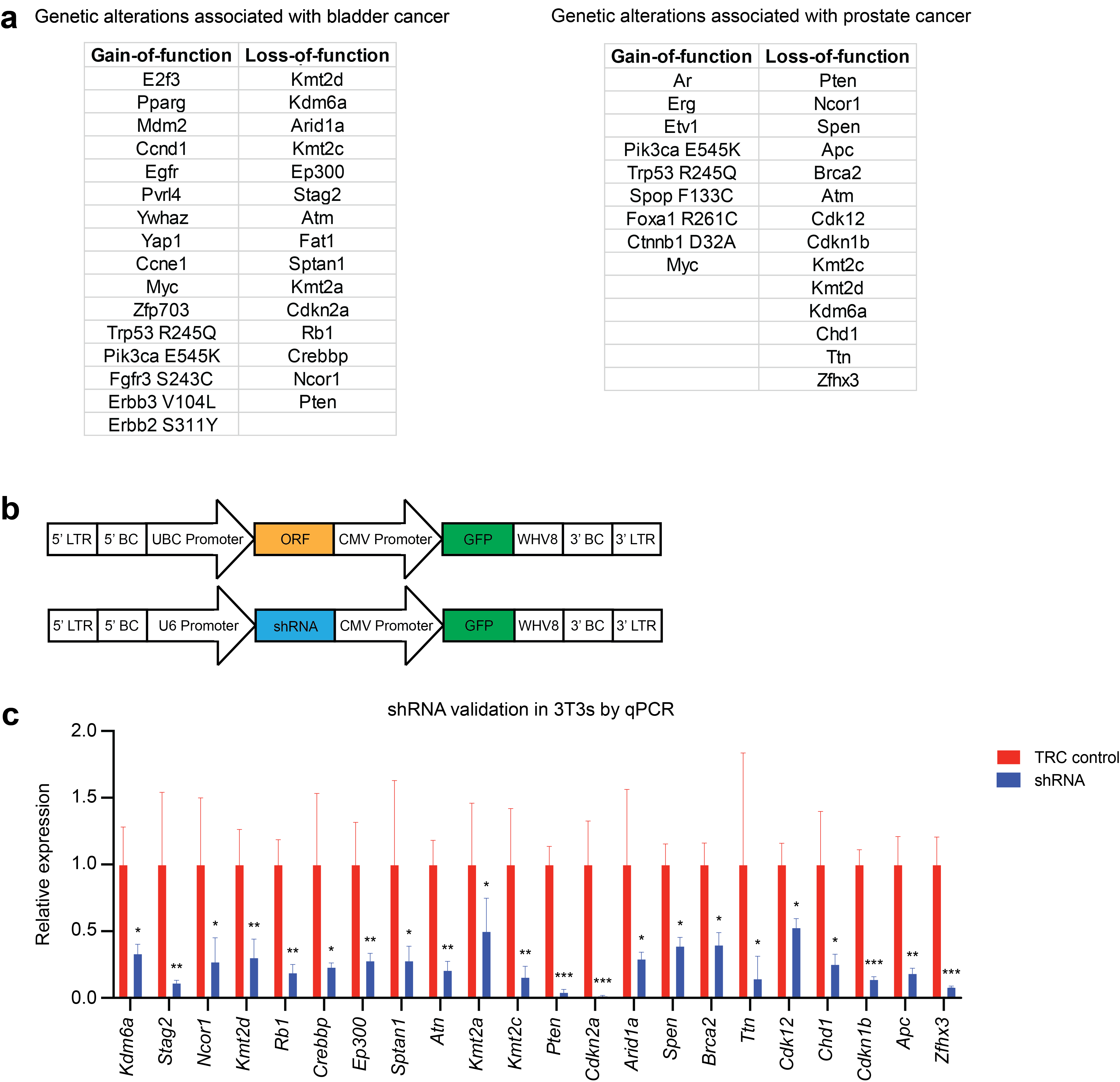

**Extended Data Fig. 2. Recurrent genetic alterations associated with bladder and prostate cancer encoded in barcoded lentiviral libraries.** (**a**) Tables showing gain-of-function and loss-of-function genetic alterations associated with bladder and prostate cancer selected for representation in cancer-specific barcoded lentiviral libraries. (**b**) Schematics of barcoded lentiviral vectors expressing open reading frames (ORF) or short-hairpin RNA (shRNA). LTR=long terminal repeat; BC=barcode; UBC=Ubiquitin C; CMV=cytomegalovirus; GFP=green fluorescent protein; WHV8=Woodchuck hepatitis virus 8 post-transcriptional regulatory element. (**c**) Plot showing relative expression of target genes as determined by quantitative polymerase chain reaction (qPCR) in 3T3 cells 72 hours after lentiviral transduction with pLKO.1-TRC control or pLKO.1 expressing select shRNA previously screened and selected for inclusion in the barcoded lentiviral libraries based on the extent of gene knockdown. qPCR reactions were performed in quadruplicate. * p<0.05; ** p<0.01, *** p<0.001.

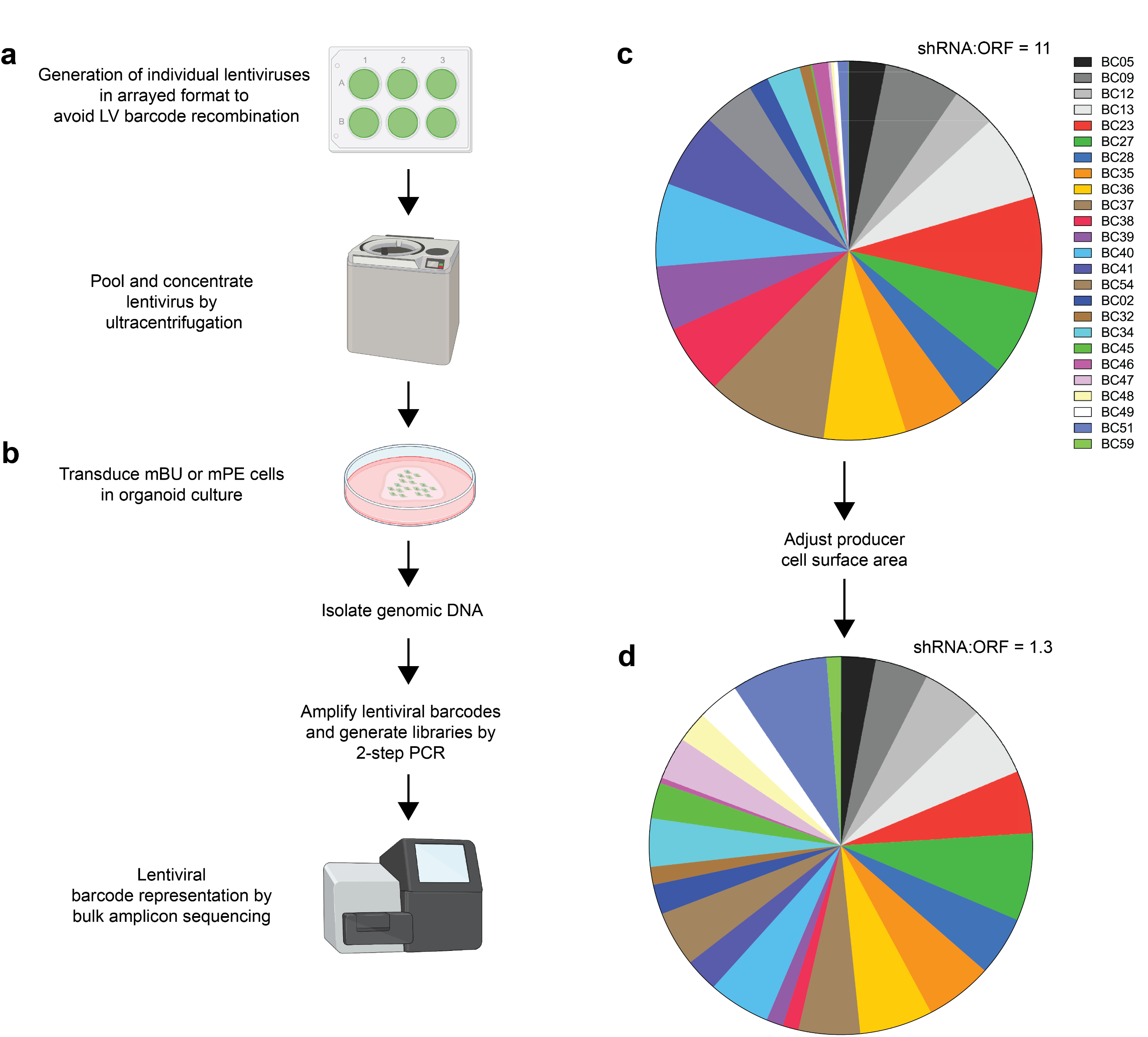

**Extended Data Fig. 3. Generation of barcoded lentiviral libraries and normalization of library representation.** Schema showing the (**a**) generation of individual lentiviruses from the library in arrayed format with subsequent pooling and concentration by ultracentrifugation and (**b**) transduction of respective mouse bladder urothelial (mBU) or prostate epithelial (mPE) cells in organoid culture with concentrated lentiviral libraries to determine lentiviral barcode representation by bulk amplicon sequencing of genomic DNA. (**c**) Representative distribution of barcoded lentiviruses within a library with skewed enrichment of shRNA relative to ORF lentiviruses. (**d**) Representative distribution of barcoded lentiviruses within a library after applying information from **c** to adjust producer cell surface area in **a** for the generation of the lentiviral library.

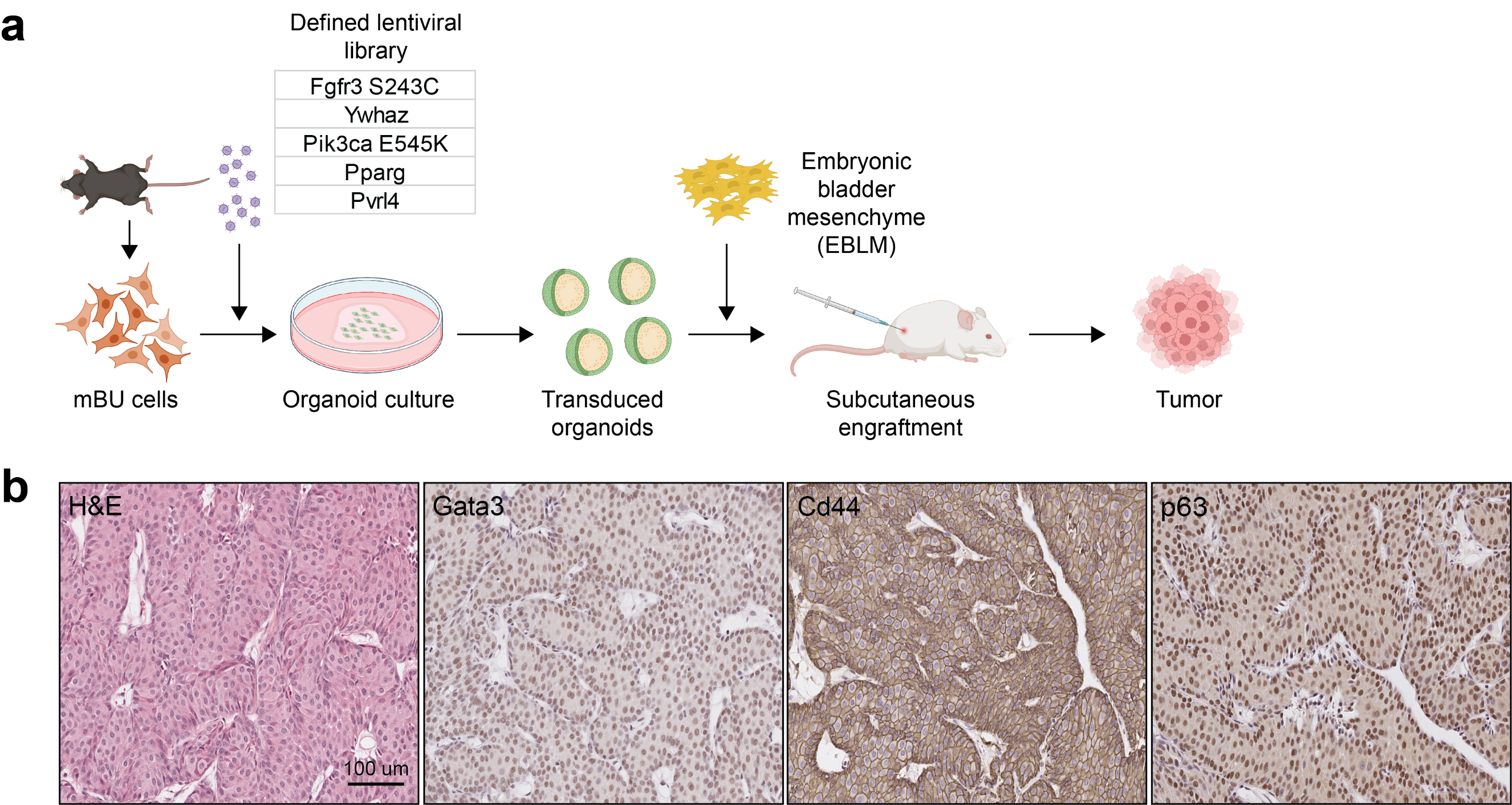

**Extended Data Fig. 4. Active mutant Fgfr3 S243C cooperates with other oncogenic factors in mouse bladder urothelial cells to drive papillary urothelial carcinoma with inverted growth pattern.** (**a**) Scheme of the mBU organoid transformation assay using a defined lentiviral library to confirm functional genotype-phenotype associations. (**b**) High-magnification images of H&E- and IHC-stained sections of a resultant tumor of the experiment in **a** with histologic features consistent with papillary urothelial carcinoma with inverted growth pattern.

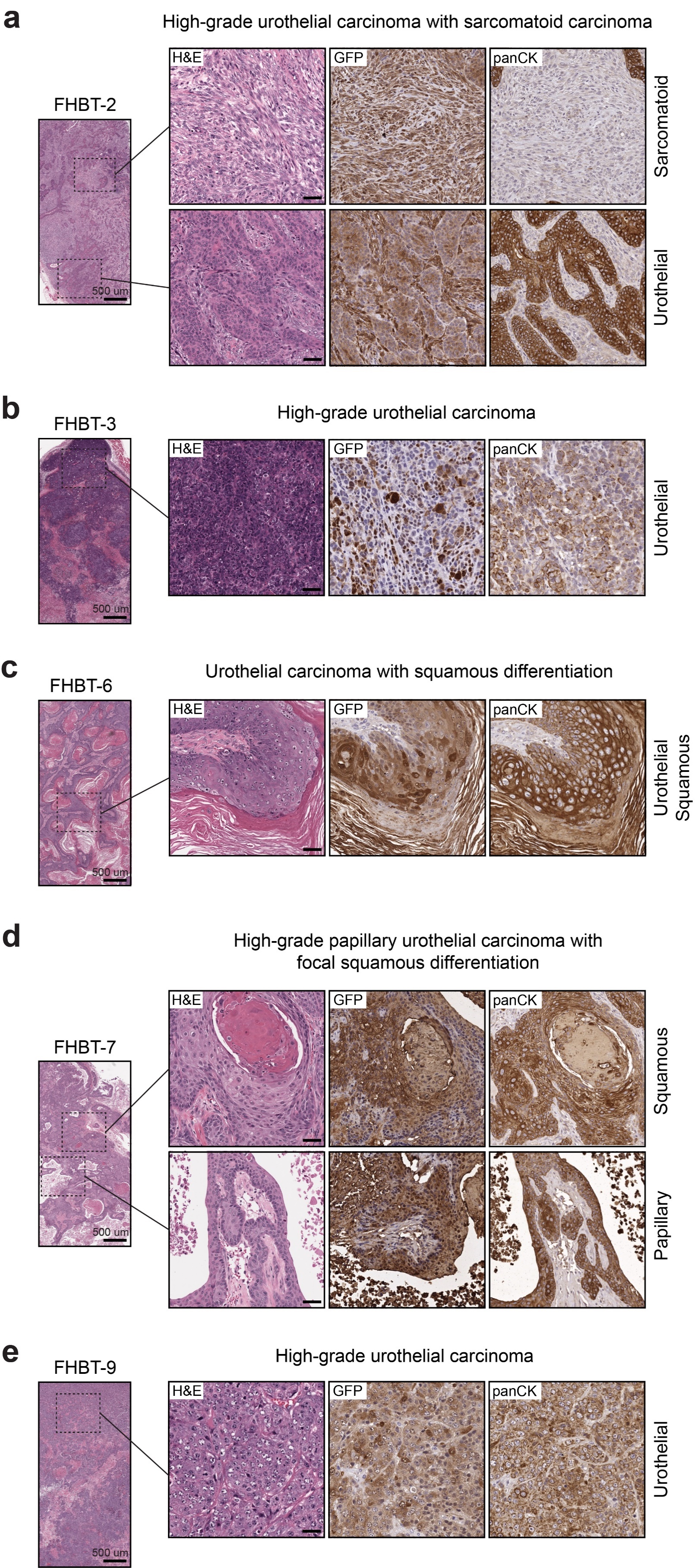

**Extended Data Fig. 5. FHBT models demonstrate diverse cancer histologies.** (**a-e**) Low- and high-magnification images of H&E-stained sections and high-magnification images of IHC-stained sections for GFP and pan-cytokeratin (panCK) expression depicting characteristic histologies. Scale bars=50 µm.

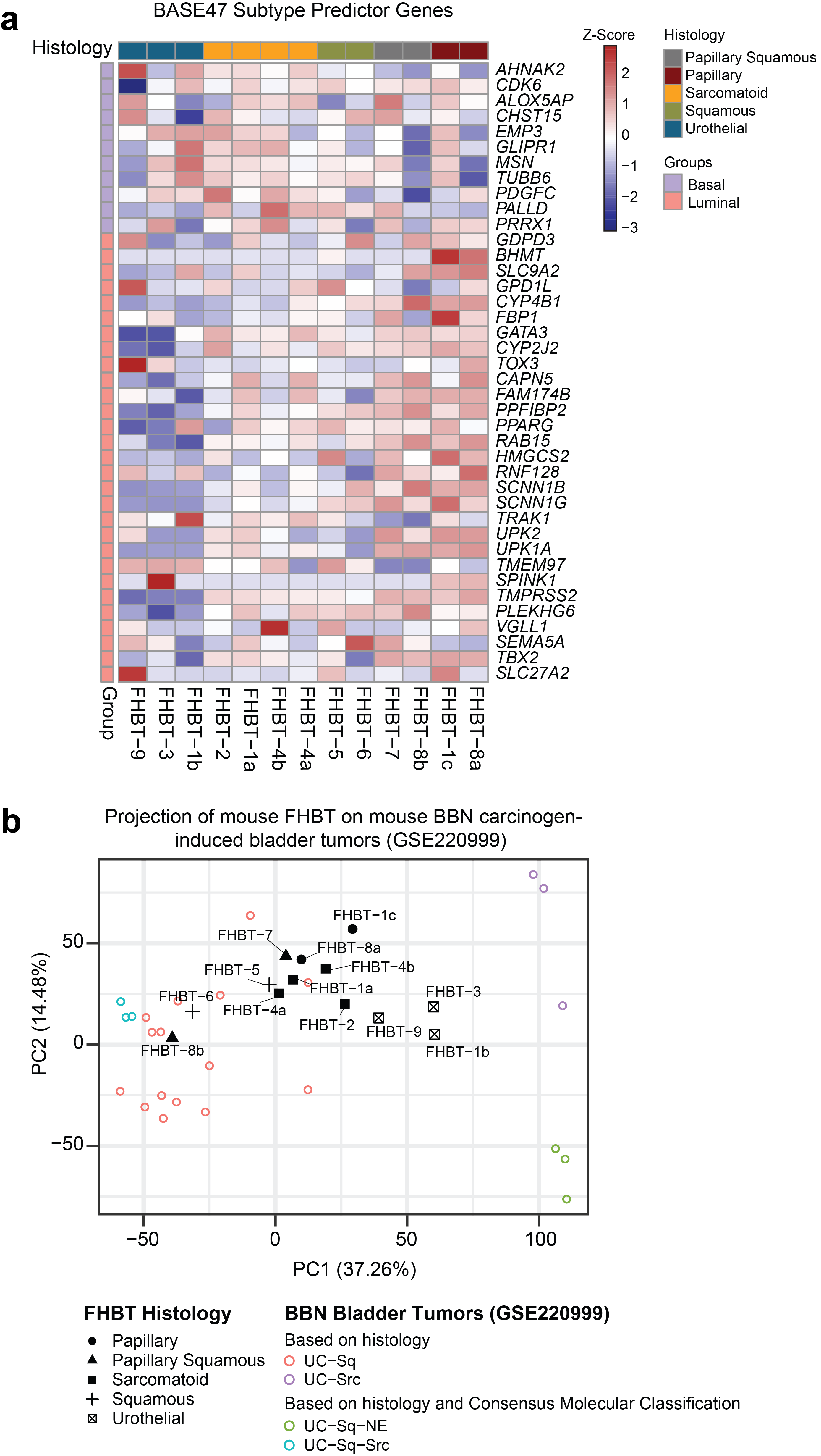

**Extended Data Fig. 6. Phenotypic diversity and relevance of FHBT models.** (**a**) Heatmap showing the histologies of the FHBT series relative to expressions of genes that constitute basal and luminal signatures for the BASE47 subtype predictor. (**b**) PCA projection plot of FHBT samples over BBN carcinogen-induced mouse bladder tumors color-coded based on histology or histology and Consensus Molecular Classification.

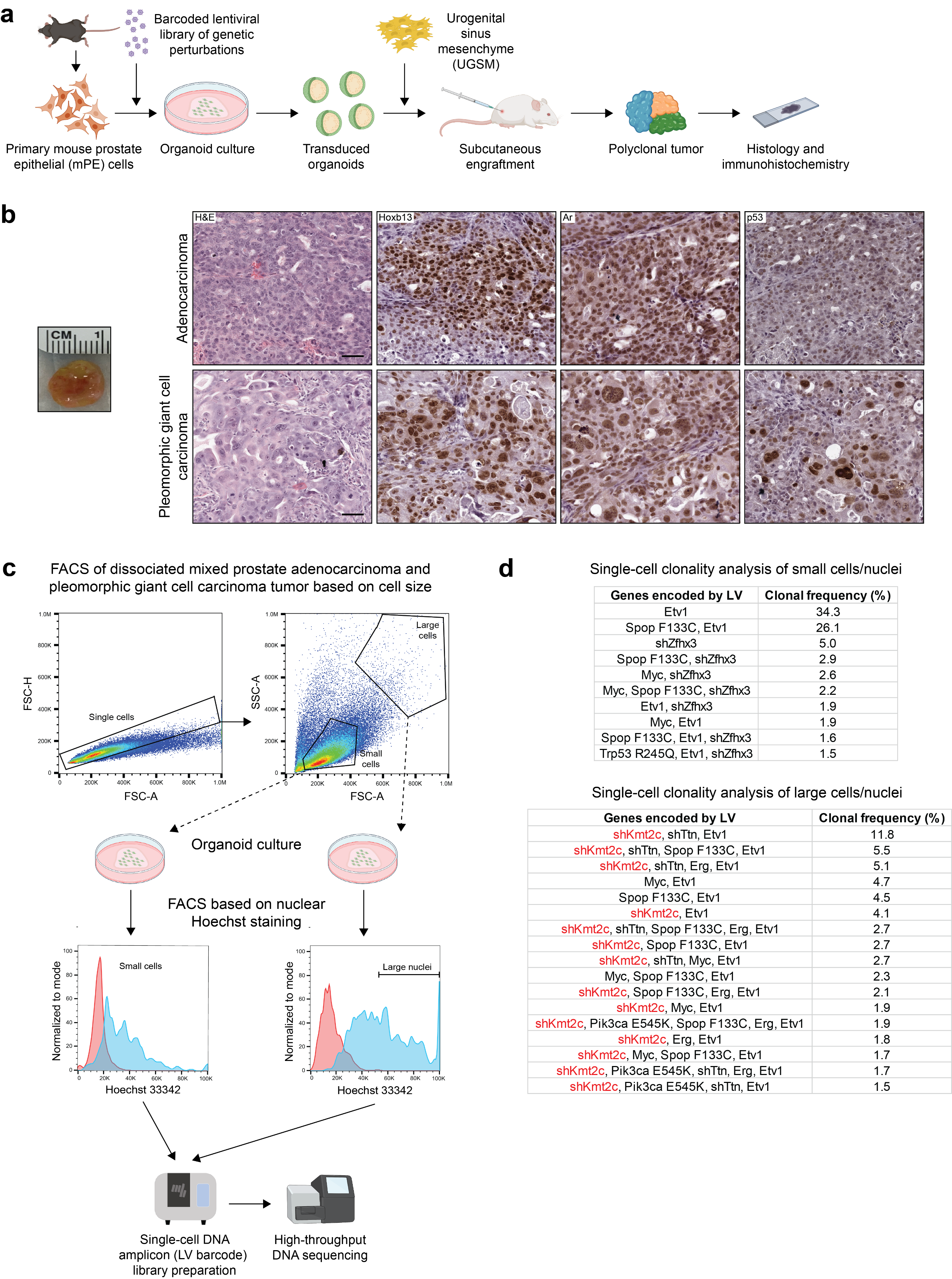

**Extended Data Fig. 7. Association of adenocarcinoma with polymorphic giant cell carcinoma of the prostate with perturbation of *Kmt2c*.** (**a**) Scheme of the mouse prostate epithelial (mPE) organoid transformation assay to uncover functional genotype-phenotype associations in prostate cancer. (**b**) *Left*, Gross image of a tumor arising from mPE transformed with PE-LVp. *Right*, high-magnification images of H&E- and IHC-stained sections of regions with high-grade adenocarcinoma and pleomorphic giant cell carcinoma. Scale bar=50 µm. (**c**) Overview of the experimental approach to enrich for prostate adenocarcinoma and pleomorphic giant cell carcinoma based on cell size and nuclear DNA content followed by single-cell lentiviral barcode enumeration. (**d**) Tables showing single-cell clonality analysis of: *Top*, tumor cells enriched for “small cells/nuclei.” *Bottom*, tumor cells enriched for “large cells/nuclei.” Highlighted in red is sh*Kmt2c* based on the enumeration of the associated lentiviral barcode.

| **Barcode (BC) #** | **Barcode sequence** | **Encoded gene** | **TRC shRNA clone ID** |
| --- | --- | --- | --- |
| 01 | GCAGATTGTA | *Yap1* | - |
| 02 | CAACGTCGAC | ORF control | - |
| 03 | TGTTGATTCA | *E2f3* | - |
| 04 | TCGCACACAC | *Pparg* | - |
| 05 | TAGTTGGCTT | sh*Kdm6a* | TRCN0000331919 |
| 06 | GGGATCGATC | *Mdm2* | - |
| 07 | AGCGTGTACA | *Ccnd1* | - |
| 08 | TCGTCCTTGA | sh*Stag2* | TRCN0000295340 |
| 09 | GAGTCATGTC | sh*Ncor1* | TRCN0000350169 |
| 10 | GGCCATTCAG | *Pvrl4* | - |
| 11 | ATCTACTCGC | *Ywhaz* | - |
| 12 | CAAACTACGT | sh*Kmt2d* | TRCN0000239233 |
| 13 | ACGATATTAG | sh*Rb1* | TRCN0000235830 |
| 14 | CAAACAATGA | sh*Crebbp* | TRCN0000012725 |
| 15 | TCGGGACAGA | sh*Ep300* | TRCN0000071207 |
| 16 | ACCGTTAGAG | *Ccne1* | - |
| 18 | ACCCACATGC | sh*Sptan1* | TRCN0000090595 |
| 20 | CTTTGACTAT | *Zfp703* | - |
| 22 | GGTCAAATCG | *Fgfr3* S243C | - |
| 23 | CCCAAATGAT | sh*Atm* | TRCN0000360328 |
| 24 | AGGCCTATCG | *Myc* | - |
| 25 | AGACTCGATG | *Erbb2* S311Y | - |
| 26 | CTATGCGTCC | sh*Kmt2a* | TRCN0000034426 |
| 27 | TCGTACGGTT | sh*Kmt2c* | TRCN0000238934 |
| 28 | GCTGGGAGTA | sh*Pten* | TRCN0000355842 |
| 29 | GAGTTCTCTA | *Egfr* | - |
| 30 | CCTATGAGTT | sh*Cdkn2a* | TRCN0000231227 |
| 31 | CCAATCTGGG | sh*Arid1a* | TRCN0000071396 |
| 32 | GAACTGAACT | TRC control | - |
| 33 | TGCATCGTTT | *Erbb3* V104L | - |
| 34 | GAAATACCTG | *Pik3ca* E545K | - |
| 35 | TTTGTTAGGT | sh*Spen* | TRCN0000226288 |
| 36 | AAACCATACT | sh*Brca2* | TRCN0000349771 |
| 37 | AACGTGTCGT | sh*Ttn* | TRCN0000362964 |
| 38 | CGCATAAGCA | sh*Cdk12* | TRCN0000361778 |
| 39 | GGTCGCAATT | shChd1 | TRCN0000096526 |
| 40 | GGTGGAACTG | sh*Cdkn1b* | TRCN0000294885 |
| 41 | TGGGTCAGCA | sh*Apc* | TRCN0000244294 |
| 43 | ATGCTGTAGG | *Prex2* | - |
| 44 | TTAAGCGCCT | *Rspo2* | - |
| 45 | ATCGGTTCAC | *Foxa1* R261C | - |
| 46 | CATGTGGACC | *Spop* F133C | - |
| 47 | TGCATGTCCT | *Ctnnb* D32A | - |
| 48 | CATTACCAAG | *Erg* | - |
| 49 | GGTCACCACA | *Ar* | - |
| 51 | ATAGAAACTC | *Etv1* | - |
| 52 | AAATGGGTAT | *Braf* G469A | - |
| 54 | CATGTCAGTC | sh*Zfhx3* | TRCN0000075411 |
| 59 | GCGTGGTTCT | *Trp53* R245Q | - |

**Extended Data Table 1. Summary of barcoded lentiviral ORF and shRNA constructs.**

| **Target gene** | **Primer 1 (5’ - 3’)** | **Primer 2 (5’ - 3’)** |
| --- | --- | --- |
| Kdm6a | ACCTAGTCCTCAGATCATACCA | GCTACTATTAGACAGGCCGTT |
| Stag2 | GGCAGATAAATTTAACCGGCTTC | GCATTATGAAAAGCAGTGATCCTC |
| Ncor1 | ATCCAGCTATGCCCTTTCAC | TGTCTGCCTTGTATTCTCCATT |
| Kmt2d | TCAGTGCTATCACCCGTACT | CACCTCACACACGATACACTC |
| Rb1 | CCTCAGCCTTCCATACTCAG | CGGAGATATGCTAGACGGTACA |
| Crebbp | TCACAATCAACATCTCCTTCCC | TGTCGATAGAGTGCTTCTAGAGT |
| Ep300 | CTCAGAAACTGTATGCCACCA | GCATCTCTACCGTCCATCAG |
| Sptan1 | GGAGGTGTATGGTGCGATG | TTGATGGAGTTGAAGGTAGCC |
| Atm | GTCACAAAGAACCATGCTTGC | CACCTTCGCAACCTCAAGA |
| Kmt2a | GAAGATGCCTGGAAGTCACT | TGCTCAATCAGAAACACAACG |
| Kmt2c | AGAAGGATGAAGAGGAAAAGCA | GGTGGTGTAGGAGGAAGAGAG |
| Pten | CACTGCTGTTTCACAAGATGATG | TTCACCTTTAGCTGGCAGAC |
| Cdkn2a | GTGCGATATTTGCGTTCCG | CTCTGCTCTTGGGATTGGC |
| Arid1a | CCTCTATCGCCTCTATGTGTCT | GCACTGCTTGATGTACCCA |
| Spen | CGACTACTTACCACGACCTTC | CACACACTAGCGATATCACAGT |
| Brca2 | GATGCCTAAACCCAGAAAGAGT | TGTGTCATCCCTCTCCAGTATC |
| Ttn | CAATGGATCTGGACAAGCGA | CACTCTCACTTGGAGTCTCAC |
| Cdk12 | AACAGACCCTACAGAGTGACT | TCGACGTTTCTTACTCCACAA |
| Chd1 | CGCCCAGCTTCATCTAATAGTG | CATCATTGTGCTTCTTCCTCTTG |
| Cdkn1b | GAGCAGACGCCCAAGAAG | GCAGTGATGTATCTAATAAACAAGGA |
| Apc | AGAATGAAGGTCAAGGAGTGG | TACTAGAACTCAAAACACTGGCT |
| Actb | GATTACTGCTCTGGCTCCTAG | GACTCATCGTACTCCTGCTTG |
| Zfhx3 | AACAACAAGATCCACCTCCAG | ACTAGGCATAACCATCTCAGGA |
| Ubc | AACATCCAGAAAGAGTCCACC | CATTCTCTATGGTGTCACTGGG |

**Extended Data Table 2. Primers used for qPCR studies to quantify relative expression of target genes.**

| **Name** | **Primer sequence (5’ - 3’)** |
| --- | --- |
| i7_7005 | CAAGCAGAAGACGGCATACGAGATGTGAATATGTCTCGTGGGCTCGGAGATGTG |
| i7_7006 | CAAGCAGAAGACGGCATACGAGATACAGGCGCGTCTCGTGGGCTCGGAGATGTG |
| i7_7007 | CAAGCAGAAGACGGCATACGAGATCATAGAGTGTCTCGTGGGCTCGGAGATGTG |
| i7_7008 | CAAGCAGAAGACGGCATACGAGATTGCGAGACGTCTCGTGGGCTCGGAGATGTG |
| i7_7015 | CAAGCAGAAGACGGCATACGAGATTCTCTACTGTCTCGTGGGCTCGGAGATGTG |
| i7_7016 | CAAGCAGAAGACGGCATACGAGATCTCTCGTCGTCTCGTGGGCTCGGAGATGTG |
| i7_7017 | CAAGCAGAAGACGGCATACGAGATCCAAGTCTGTCTCGTGGGCTCGGAGATGTG |
| i7_7018 | CAAGCAGAAGACGGCATACGAGATTTGGACTCGTCTCGTGGGCTCGGAGATGTG |
| i7_7023 | CAAGCAGAAGACGGCATACGAGATGCAGAATTGTCTCGTGGGCTCGGAGATGTG |
| i7_7024 | CAAGCAGAAGACGGCATACGAGATAACCGCGGGTCTCGTGGGCTCGGAGATGTG |
| i7_7025 | CAAGCAGAAGACGGCATACGAGATACTAAGATGTCTCGTGGGCTCGGAGATGTG |
| i7_7026 | CAAGCAGAAGACGGCATACGAGATGTCGGAGCGTCTCGTGGGCTCGGAGATGTG |
| i5_5001 | AATGATACGGCGACCACCGAGATCTACACAGCGCTAGTCGTCGGCAGCGTCAGATGTGT |
| i5_5002 | AATGATACGGCGACCACCGAGATCTACACGATATCGATCGTCGGCAGCGTCAGATGTGT |
| i5_5007 | AATGATACGGCGACCACCGAGATCTACACACATAGCGTCGTCGGCAGCGTCAGATGTGT |
| i5_5008 | AATGATACGGCGACCACCGAGATCTACACGTGCGATATCGTCGGCAGCGTCAGATGTGT |
| i5_5009 | AATGATACGGCGACCACCGAGATCTACACCCAACAGATCGTCGGCAGCGTCAGATGTGT |
| i5_5010 | AATGATACGGCGACCACCGAGATCTACACTTGGTGAGTCGTCGGCAGCGTCAGATGTGT |
| i5_5013 | AATGATACGGCGACCACCGAGATCTACACAACCGCGGTCGTCGGCAGCGTCAGATGTGT |
| i5_5014 | AATGATACGGCGACCACCGAGATCTACACGGTTATAATCGTCGGCAGCGTCAGATGTGT |

**Extended Data Table 3. Primers used for 2º PCR to incorporate dual-indexed Illumina sequencing adaptors for bulk amplicon sequencing.**
